# Motor cortical beta transients delay movement initiation and track errors

**DOI:** 10.1101/384370

**Authors:** Simon Little, James Bonaiuto, Gareth Barnes, Sven Bestmann

## Abstract

Motor cortical activity in the beta range (13-30 Hz) is a hallmark signature of healthy and pathological movement, but its behavioural relevance remains unclear. Recent work in primates and human sensory cortex suggests that sustained oscillatory beta activity observed on average, may arise from the summation of underlying short-lasting, high-amplitude bursts of activity. Classical human movement-related event-related beta desynchronisation (ERD) and synchronization (ERS) may thus provide insufficient, non-dynamic, summaries of underlying focal spatio-temporal burst activity, limiting insight into their functional role during healthy and pathological movement.

Here we directly investigate this transient beta burst activity and its putative behavioural relevance for movement control, using high-precision magnetoencephalography (MEG). We quantified the subject-specific (n=8), trial-wise (n>12,000) dynamics of beta bursts, before and after movement. We show that beta activity on individual trials is dominated by high amplitude, short lasting bursts. While average beta changes generally manifest as bilaterally distributed activity (FWHM = 25mm), individual bursts are spatially more focal (FWHM = 6 mm), sporadic (1.3 −1.5/s), and transient (mean: 96 ms).

Prior to movement (the period of the classical ERD), the timing of the last pre-movement burst predicts movement onset, suggesting a role in the specification of the goal of movement. After movement (the period of the classical ERS), the first beta burst is delayed by ~100ms after a response error occurs, intimating a role in error monitoring and evaluation.

Movement-related beta activity is therefore dominated by a spatially dispersed summation of short lasting, sporadic and focal bursts. Movement-related beta bursts coordinate the retrieval and updating of movement goals in the pre- and post-movement periods, respectively.

## Introduction

Cortical activity in the beta frequency range (13-30 Hz) has been recognised for nearly a century, and occurs with systematic activity changes before, during, and after movement (Cassim et al., 2000; Fetz, 2013; Houweling et al., 2010; Hummel et al., 2003; Manganotti et al., 1998; Müller et al., 2003; Pfurtscheller, 1981; Pfurtscheller and Lopes, 1999; Ray and Maunsell, 2015, 2010; Schulz et al., 2011; Shadlen and Newsome, 1998, 1994, Stančák et al., 1997, 2000; Toma et al., 2002).

Pre-movement beta activity is, on average, characterized by slow and spatially diffuse decreases in beta activity beginning up to a few seconds prior to movement with a minimum at movement onset (van Wijk et al., 2012). This average pre-movement beta signal is influenced by a wide range of processes, including general preparatory processes (Tzagarakis et al., 2010), decisions for actions (Doyle et al., 2005; Gould et al., 2012; van Wijk et al., 2009), movement kinematics (Houweling et al., 2010; Hummel et al., 2003; Manganotti et al., 1998; Stančák et al., 1997; Toma et al., 2002) and inhibition of premature and unwanted responses (Khanna and Carmena, 2017; Wessel and Aron, 2017). Following movement, beta activity increases before gradually returning to baseline, and this increase has been linked to processes relating to updating of movement outcomes (Fine et al., 2017; Tan et al., 2014, 2016; Torrecillos et al., 2015).

In addition to healthy movement control, understanding the functional role of cortical beta activity appears vital to unlocking understanding of the pathophysiology of diseases of the human motor system (Hammond et al., 2007; Heinrichs-Graham et al., 2013; de Hemptinne et al., 2013; Meziane et al., 2015; Teodoro et al., 2017). However, despite such a reliable hallmark neural signature of healthy and pathological movement, and its robust relationship to a variety of broader processes including top-down communication (Bressler and Richter, 2015; Buschman and Miller, 2007), status-quo maintenance (Engel and Fries, 2010), sensori-motor integration (Androulidakis et al., 2006, 2007; Baker, 2007; Gilbertson et al., 2005), and attention (Saleh et al., 2010), a unifying theory for the behavioural role of movement-related beta activity remains elusive.

Recent animal work and isolated reports in humans suggests that slow, sustained, changes in beta activity pre- and post-movement may not sufficiently summarize the trial-wise dynamics in beta amplitude (Feingold et al., 2015; Leventhal et al., 2012; Murthy and Fetz, 1992, 1996; Pfurtscheller et al., 1996). Specifically, work in the frontal cortex of non-human primates suggests that cortical beta activity is characterised by transient bursting that may only appear to be temporally sustained if averaged over multiple trials (Feingold et al., 2015; Lundqvist et al., 2016). Instead, movement-related beta activity may be dominated by infrequent, non-rhythmic bursts, with the highest probability of occurrence following movement, but with a wide temporal dispersion (Feingold et al., 2015). This notion of short lasting beta bursts in humans is supported by recent work on the putative functional role of beta transients in sensory processing. In somatosensory cortex, beta bursting relates to tactile sensory discrimination, whereby non-detection is linked to higher burst rate and temporal coincidence with sensory input (Shin et al., 2017). Trial-averaged beta activity may therefore reflect the underlying cortical burst rate, which will itself stochastically determine the probability of the very largest bursts occurring across trials at behaviourally relevant time points (Jones et al., 2009; Sherman et al., 2016; Shin et al., 2017). Burst occurrence may be linked to transient inhibitory activity directed to sensory processing, an idea supported by biophysical modelling of inter-laminar neural dynamics (Shin et al., 2017). However, whether beta has an active processing and information transmission role which may generalise to other brain systems (e.g. motor) is unknown (van Ede et al., 2018). Importantly, the transient, infrequent and non-rhythmic dynamics of beta bursts are at odds with the slowly changing and spatially diffuse average signals that have formed the basis for most accounts of the functional role of movement-related beta activity in health and disease (Cheyne, 2013; Engel and Fries, 2010; Kilavik et al., 2012; van Wijk et al., 2012).

Specifically, it would seem difficult to reconcile a role for beta in sustained facilitation of the current motor state if bursts are sporadic, short-lasting events which do not reliably time-lock to behaviour and dominate the changes in beta power both within and across trials. Here we address whether burst characteristics such as rate, size and timing might be primary drivers of behaviour.

To investigate the role of cortical bursting activity in healthy subjects before and after movement, we analyse trial-wise dynamic beta burst activity from motor cortex as obtained from a high precision MEG dataset using subject specific head-casts and detailed cortical models (Bonaiuto et al., 2017; Meyer et al., 2017; Troebinger et al., 2014a). We investigate the precise relationship between the distribution of beta bursts and the spatial and temporal profile of the classical event-related desynchronisation and synchronisation (ERD and ERS). Furthermore, we examine the specific behavioural relationship between the rate and timing of beta bursts to movement initiation and error monitoring.

## Results

We recorded high signal-to-noise (SNR) MEG, using individual head-casts (Bonaiuto et al., 2018; Meyer et al., 2017; Troebinger et al., 2014b, 2014a), in 8 healthy subjects during a probabilistically cued movement selection task (Fig.1 A,B). This provided us with pre- and post-movement measures of beta activity, with over 12,000 trials (per subject M=1620, SD=763.7)(Bonaiuto et al., 2017). Multiple runs (1-4) were recorded per subject over several days and within each recording of 15 minutes, head movements were only 0.23 ± 0.04, 0.25 ± 0.05 and 0.99 ± 0.54 mm in the x, y and z directions respectively (Fig.1 C), supporting high SNR individual datasets.

**Figure 1.**
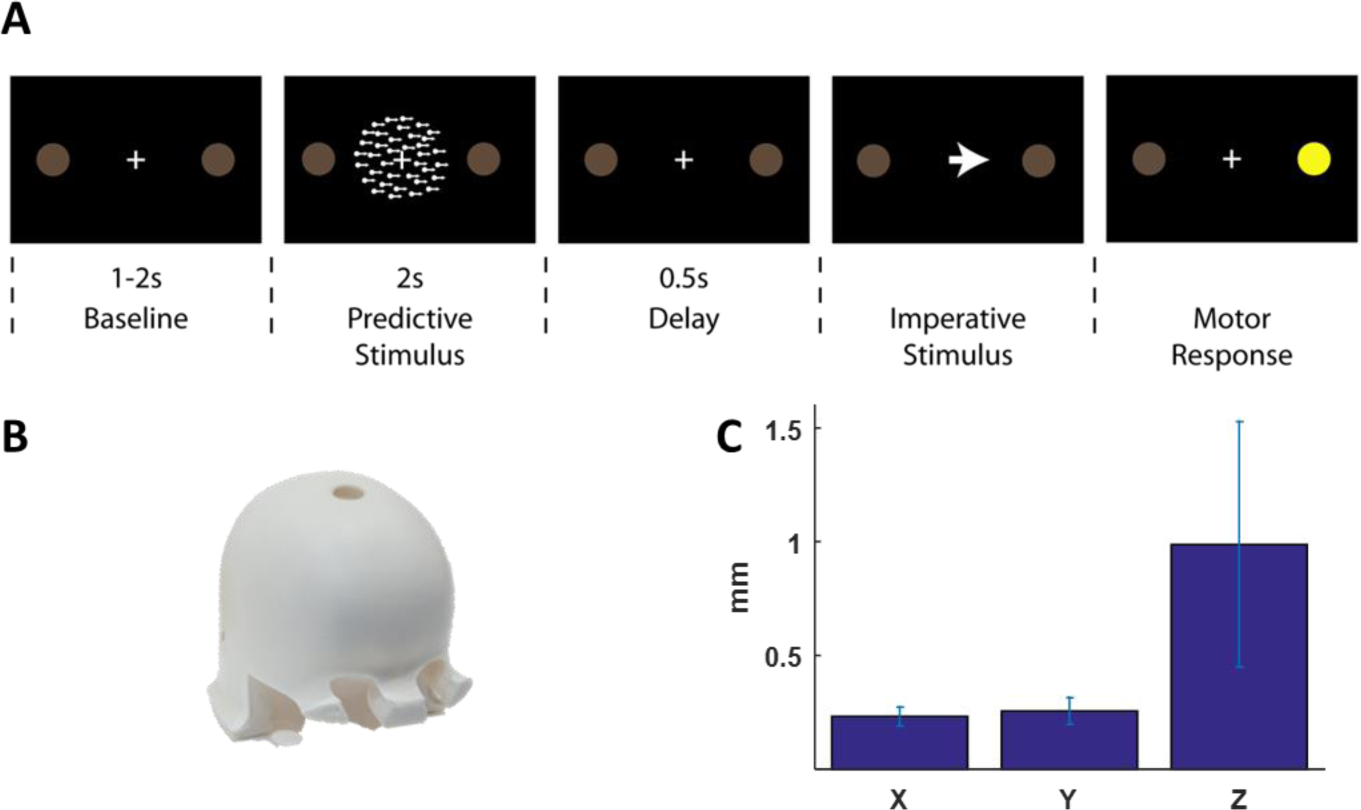
Task design and high-precision, head-cast, MEG set-up. **A** Behavioural paradigm with baseline period followed by a random dot kinetogram (RDK) indicating the likely direction of a subsequent (congruent or incongruent) instruction cue, signalling a left or right button response. **B** Subject-specific head-casts improve within and between session co-registration accuracy and reduce within-run head movements, facilitating the acquisition of large, high SNR, datasets. **C** Mean head movements across all subjects (± SEM errorbars) are on average <1mm.

Source inversion and virtual electrode creation at the contralateral primary motor cortex (see methods) disclosed a broadband (2-100 Hz) time series for each trial. Frequency specific changes in movement-related activity were first examined using a time varying spectrogram (averaged across all trials), shown here for a representative subject (Fig 2A; top panel). The spectrogram illustrates the classical, average (across trials), reduction in beta activity (13-30 Hz) prior to movement (ERD - blue), and, average increase after movement (ERS – red); also shown specifically for the beta band alone (Fig 2A; middle panel).

**Figure 2.**
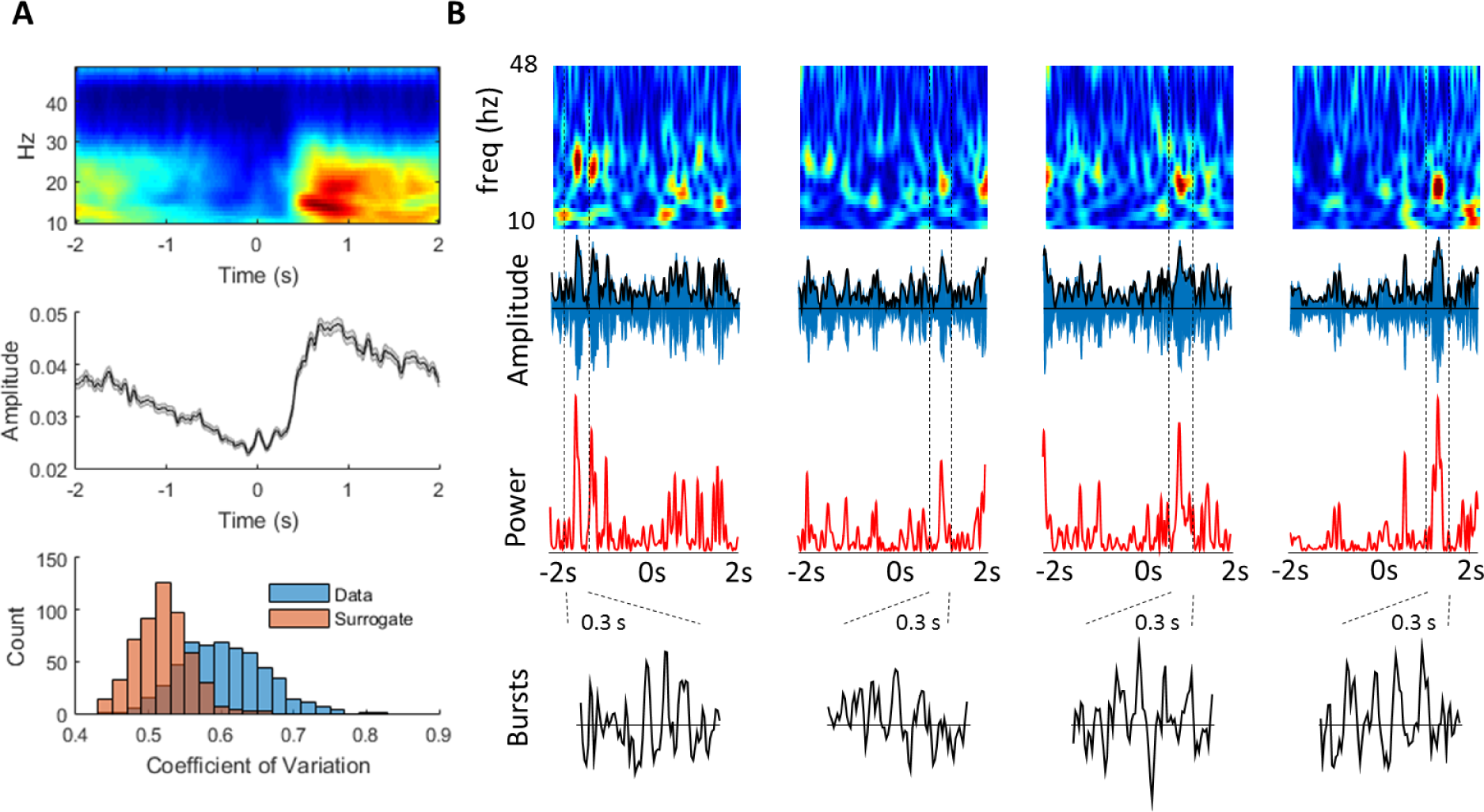
Individual trial data contains highly dynamic beta signal. **A** Single subject, average time-frequency spectrogram (wavelet decomposition; see methods) showing a reduction in spectral activity, specific to the beta band, in the period prior to movement, followed by an increase following movement (subject 2; top panel). Middle panel: time series of average beta (13-30 Hz) ERD and ERS. Bottom panel: Distribution of beta amplitude variability (coefficient of variation (CV)) across trials, compared to a phase-shuffled, spectrally matched, surrogate dataset from the same subject. The right shift of the CV demonstrates higher beta amplitude variability in real data compared to the shuffled, surrogate data set. **B** Four consecutive individual trials from the same subject (trials 250-254), shown as amplitude spectrograms (top), beta filtered times series (blue) with superimposed amplitude envelope (black; second row panels), power (amplitude.^2; red, third row panels) and 4 individual, magnified, unfiltered, burst time series (black; bottom panel; periods taken between dashed vertical black lines; 300 ms duration).

This gradually modulating signal has been linked to mechanisms supporting the current motor state as well as inhibitory processes relating to upcoming movement planning and preparation (Engel and Fries, 2010). Proposals about the functional relevance of movement-related beta activity generally rest on the implicit assumption that individual trials show a similar pattern of activity to the average ERD and ERS signal, and that the behaviourally relevant component is therefore the *slowly* modulating magnitude of the beta signal. However, we next show that the coefficient of variation of the beta amplitude is consistently higher than expected when compared to a phase-shuffled surrogate data, which is spectrally identical to that of the representative subject (Fig 2A, lower panel; see methods). Subsequently, individual trials from this same representative subject are plotted as spectrograms, filtered signal (with amplitude enveloped superimposed), beta power (amplitude .^2), and individual selected beta burst time series. This reveals the highly dynamic activity in the un-averaged data (Fig 2B) which does not obviously accord with the average and gradual changes in beta shown in the average spectrogram (Fig 2A). Conventionally, beta amplitude envelopes have been averaged over trials to remove and smooth out such dynamic fluctuations, with attention focused (and behavioural correlations made) on the slowly changing, mean signals that results. We here asked whether such dynamic changes in beta activity contributed to classical average signals, and whether these transient bursts had significant behavioural relevance. We use “ERD” and “ERS” to denote average (across trials) changes in the magnitude of the beta signal before and after movement, respectively. At the single trial (non-averaged) level we use “amplitude” to refer to the time varying envelope of the beta signal (√power; Fig B second panels, black line).

In order to formally assess the relationship between the average beta ERD, ERS and transient burst activity, we empirically determined in each individual and separately for the ERD and ERS period a threshold for defining burst activity. These thresholds were quantified in terms of standard deviations (SD) of the beta amplitude variability over time, above the median beta amplitude for the pre- (−3000ms → 0ms) and post- (0 → 2000ms) movement periods. In order to define these thresholds, we used the method of Shin et al. (Shin et al., 2017), and for each subject we correlated the trial-wise beta amplitude with beta burst count across a range of different thresholds for the pre- and post-movement periods separately. The burst definition threshold, across all subjects, that resulted in the highest correlation between burst rate and mean (within trial) beta amplitude, was found to be 1.75 SDs above the median for the pre-movement and post-movement periods. This threshold was then used on each subject (on instruction cue and response locked datasets) so that their relative thresholds were matched (1.75 SDs above beta amplitude median), although absolute thresholds could differ across subjects according to their data and SNR.

### Beta burst probability is modulated by movement

The temporal distribution of motor cortical beta bursts in relation to movement remains unexplored in humans. To assess whether the beta ERD and ERS might contain significant beta burst activity, we examined the relationship between the burst probability and the ERD and ERS in primary motor cortex during the pre- and post-movement periods, respectively (Fig 3A).

**Figure. 3.**
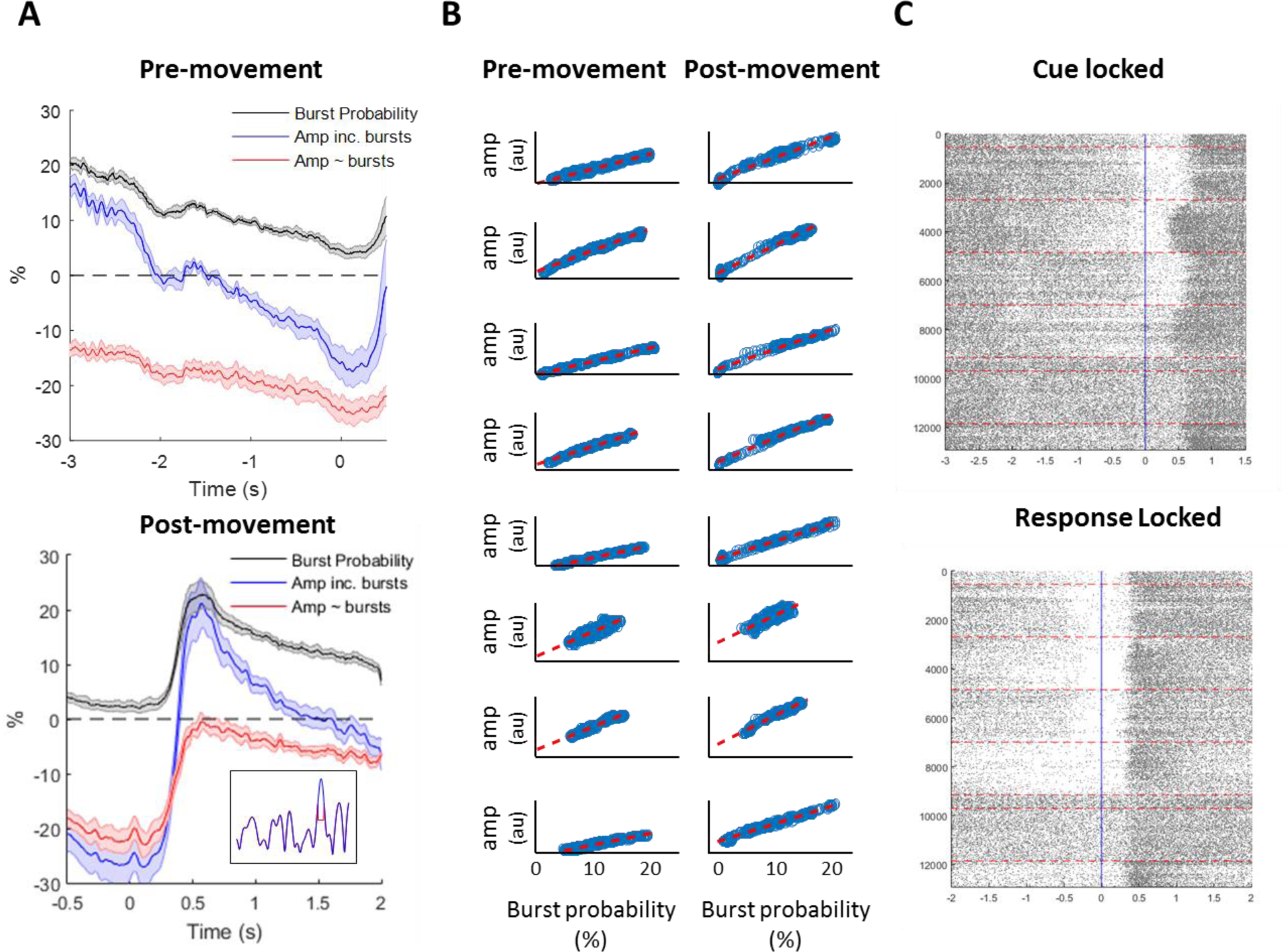
Burst probability is modulated by movement and accounts for the classical ERD and ERS. **A** Normalised (% versus pre-movement mean) beta amplitude ± SEM (blue) is shown in the pre-movement period with burst probability (%) overlaid above (black). Beta ERD with bursts replaced by the mean beta amplitude is shown overlaid (red) demonstrating that replacement of low numbers of sporadic (comprising just 11.9% of total pre-movement time) bursts markedly flattens the ERD (top panel). Below we show the same analysis for the post-movement period. Note that the burst probability accurately tracks the classical change in the beta amplitude and also that replacement of short periods of intermittent bursting (12.8% of total time series) by mean power removes the classical ERS peak and attenuates the late ERS slope. Example of a single subject / trial (inset; 0 – 1000ms post-movement) beta amplitude (blue) shown with single burst replaced with the mean (red) **B** Burst probability (across all trials) for each subjects is correlated against the normalised, time resolved ERD and ERS demonstrating a strong and consistent relationship between them. **C** Raster plot showing timing of each individual burst (single dash = peak of burst) for all 8 subjects (>12,000 trials) using the empirically defined (1.75 sd > median) threshold (individual subjects divided by dashed red lines). This highlights a notably consistent relationship of burst timing to movement onset with a gradual reduction in burst probability in the time building up to movement and then a significant increase (weakly temporally locked) after movement.

Pre-movement, the average beta burst rate was 1.3 ± 0.08 bursts per second (bps; −3s to 0s prior to instruction cue). Normalised average ERD beta (across trials) in the same pre-movement period (% relative to pre-movement mean) decreased from 16.3.3 ± 2.1 % (−3 s prior to the instruction cue presentation) to – 15.9 ± 3.2 (at the instruction cue; Fig 3C, top panel). We then calculated the time resolved burst probability over the same period (for each time point; burst probability = number of trials in which there is a burst at that time point / total number of trials; Fig 3C, black line). During the pre-movement period the beta burst probability across trials fell from 20.3 ± 0.7 % to 4.5 ± 1.0 % at the instruction cue (Fig 3C, top panel). In order to assess whether beta bursts made a significant contribution to the shape of the average (classical) ERD signal, we then mathematically replaced the short periods during each beta burst with the mean (pre-movement) beta amplitude. Therefore, if beta really were sustained oscillations at the single trial level (i.e. bursts were noise, unrelated to movement) one would expect that replacing sparse bursts would lead to only a small shift in all timepoints of the ERD downwards but not greatly change the overall shape. Alternatively, if beta bursts appreciably contribute to the ERD, we predicted that removing bursts would considerably flatten the shape of the ERD curve. This was borne out by the data, in which bursts comprised only 11.9 ± 0.2 % of the total time course (pre-movement) and yet their replacement with mean beta amplitude led to a significant reduction in early ERD from 16.3.3 ± 2.1 % to −13 ± 1.6 % (−3 seconds prior to instruction cue, t7=−10.5; p<0.001). There was a smaller reduction in the late pre-movement period from −15.9 ± 3.2 % to −24.2 ± 2.3 % (onset of instruction cue, t7=−5.4; p=0.001) with therefore an overall effect of flattening of the ERD curve.

Post-movement, the average beta burst rate was 1.51 ± 0.08 bps and the average beta ERS signal increased from −25.8 ± 3.9 (button press) to 21.2 ± 4.9% at the peak (568 ms). Over the same period, the beta burst probability increased from 2.3 ± 1.0 to 22.9 ± 2.5 %. Therefore, even at the peak of the ERS, bursts were infrequent across trials at that exact moment (< 1 in 4 trials). Furthermore, during the ERS period, rather than aligning to movement, bursts were temporally delayed and widely distributed across the post-movement period with mean onset of the maximum of the first burst at 681 ± 34 ms post button press with an average 404 ± 12.5 ms within subject standard deviation (ie variability across trials for each individual subject). We then repeated our burst (mean) replacement analysis in the post-movement period with the prediction this would specifically attenuate the classical ERS (rather than lower all time points across the time series, our alternative hypothesis if bursts were noise, unconnected to behaviour). Here, replacement of the bursts, comprising just 12.8 ± 0.32 % of the total time course, significantly attenuated the classical ERS, reducing it from a maximum of 21.2 ± 4.9 % above the mean (568 ms) to −0.6 ± 1.8 % following burst replacement. Importantly, replacing periods of higher beta amplitude with the (lower) mean beta amplitude will inevitably reduce the results (modified) ERD and ERS across trials. However, what is notable about replacing the beta bursts was that it was the shape of the ERD (gradient reduced) and ERS (peak flattened) that changed as opposed to a general (time independent) shift across the whole period. Furthermore, the complete negation of the ERS peak despite removal of just a small amount (12.8%) of the total data in the form of short lasting, infrequent (less than 1 burst in 4 trials at peak) bursts (threshold of 1.75 sd above the median captures ~ top 12% of data), strongly suggests that the cortical ERD and ERS are dominated by burst probability rather than a slowly changing drift in amplitude.

We therefore sought to directly relate the shape of the time resolved burst probability with that of the ERD and ERS by correlating the shape of these two curves with each other over time. In contrast to the method for empirically determining the threshold (correlating burst rate with beta amplitude *across trials*) we here correlate the time resolved burst probability (Fig 3A, black line) against the ERD and ERS *over time* (Fig 3C, blue line). This quantifies, over the timecourse of a trial, the relationship between the shape of classical ERD and ERS and the shape of beta burst probability during the trial (Fig 3B), separately for the pre-movement and post-movement periods and can be considered as correlating the blue and black curves (Fig 3A) against each other. At each time-point, burst probability and the ERD and ERS at that same point (across all trials) were closely related (mean R^2^ of 0.93 ± 0.02 in the pre-movement period and a mean R^2^ 0.93 ± 0.03 in the post-movement period; Fig 3B). This indicates that average beta changes (as performed in the majority of conventional analyses) is dominated by burst probability across trials (Fig. 3B). In order to directly visualise the relationship between burst timing and behaviour, we plotted the timing of beta bursts for all subjects and all trials separately for the pre- and post-movement periods as a raster plot (Fig 3C). This shows a striking consistency in the distribution of bursts across subjects, with minimal bursts around movement onset, and then a temporally diffuse increase in burst probability after movement.

### Beta bursts are larger than matched surrogate data

We next examined whether the variability of the beta signal could be differentiated from a spectrally matched, surrogate (phase-shuffled) dataset across all subjects. We found that the coefficient of variation of the beta amplitude was increased in all subjects in the real data compared to the surrogate for the pre- (t_7_=21.3; p<0.0001; Fig 4A) and post-movement periods (t_7_=26.4; p<0.0001; Fig 4A) respectively (Feingold et al., 2015; Little et al., 2012a). We then focused our attention specifically on the empirically defined bursts (data > 1.75 sd above the median). Bursts can vary in terms of amplitude as well as duration and previous evidence in primates suggests that the distribution of cortical beta amplitude is skewed towards high amplitude bursts when compared to surrogate data in primates (Feingold et al., 2015). The distribution of peak beta burst amplitudes in our dataset, across all time points, was therefore compared with the peak beta burst distribution from the same surrogate (phase shuffled) dataset. The comparison demonstrated that there were a greater number of larger bursts in the original data compared to the surrogate dataset and this was significant for all burst sizes above the 30^th^ percentile (p_FDR_<0.05, Fig. 4B). Burst duration was also longer than in the surrogate dataset, but in a restricted range (300-400 ms, p_FDR_<0.05). In fact, most bursts were of short duration with an average burst duration of 96.2 ±8.8 ms and 94.2 ± 10.4 ms in the pre- and post-movement periods, respectively (t7= 0.7,p=0.49). Burst characteristics were significantly larger in terms of height (t7=2.3; p=0.049), but showed no difference in length (t_7_=−0.7; p=0.49) w the pre- and post-movement period.

**Figure 4.**
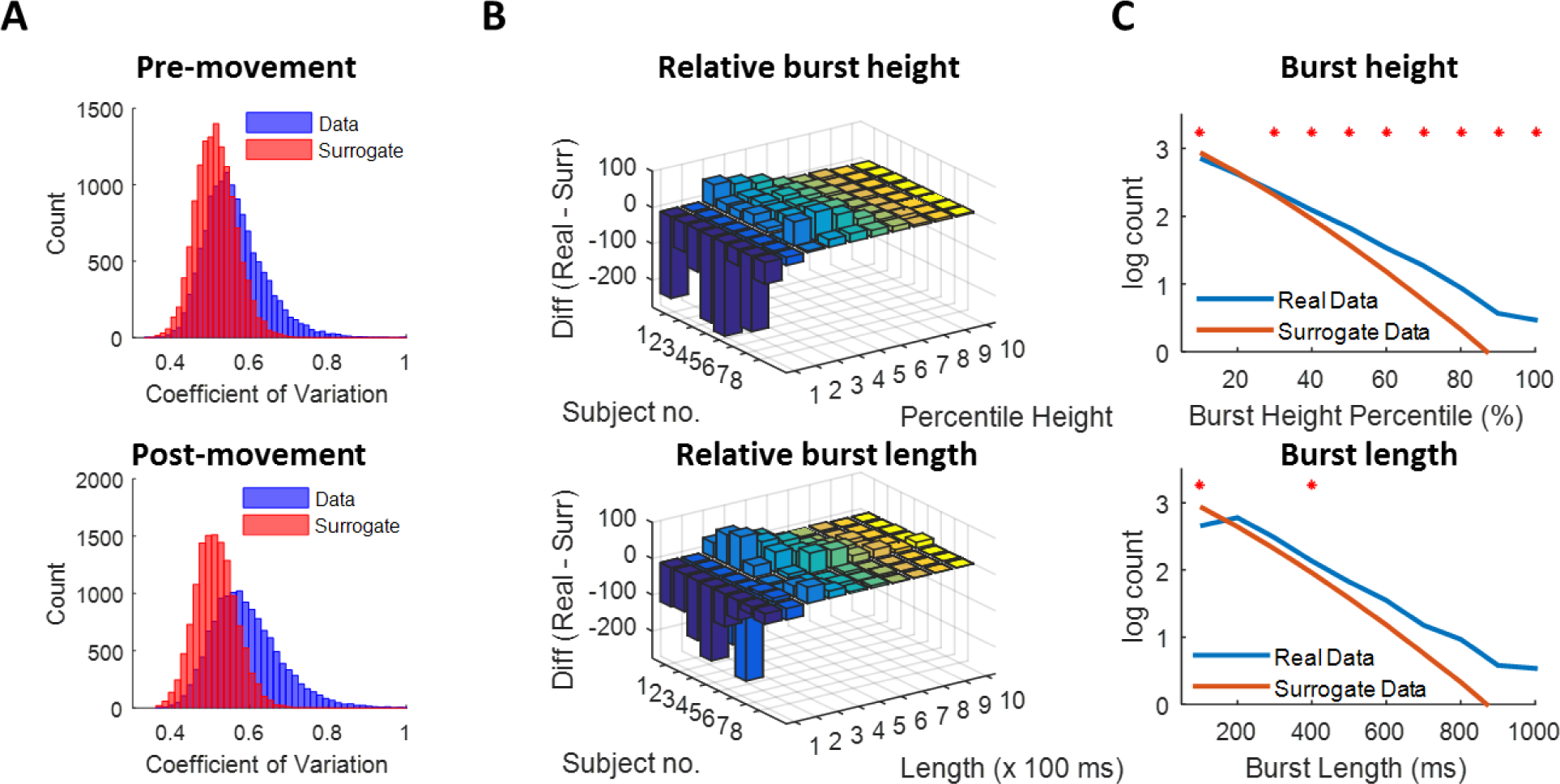
Beta amplitude shows high powered bursts. **A** Histograms of the coefficient of variation of beta amplitudes (across all trials) in real and surrogate data for the pre- (ERD) and post- (ERS) movement periods. Note that the distribution for the real data is shifted to the right of the surrogate data which was significantly increased in both the pre (ERD) and post (ERS) movement periods. **B** Difference in burst height and length counts shown for each subject compared to bursts from a surrogate phase-shuffled dataset (burst height/length real data minus burst height/length surrogate data). This demonstrates a relative reduction in the number of the smallest and shortest bursts in the real compared to surrogate data, with an increased in larger and mid-length bursts. **C** Group level line histogram of the log10 mean burst counts for different height and lengths for real versus surrogate data showing a significant difference (*, False Discovery Rate (FDR) corrected).

There was a tendency of M1 beta oscillations to produce more, higher powered and longer, bursts than would be expected from spectrally matched surrogate data (Fig. 4). Therefore, this difference in size and length, in addition to their strong modulation by behaviour (Fig. 3) further supports the contention that beta bursts might form part of active processing mechanisms rather than simply representing the noise on top of a slowly changing classical ERD and ERS signals (van Ede et al., 2018; Spitzer and Haegens, 2017).

Thus far we have formally examined the properties (height and duration) of the burst amplitude (Fig 4 A,B). We next proceeded to examine individual bursts in the time series domain to investigate their consistency with regards to shape in the pre- and post-movement periods and within and across subjects (Sherman et al., 2016; Shin et al., 2017). For each subject therefore we collected all the bursts (unfiltered) together from the pre- and post-movements periods separately, aligned to the maximum of the burst amplitude peak (± 100ms). We then averaged across these bursts to derive the mean burst shape (Fig 5A; average burst for each subject shown). In order to quantify the homogeneity of bursts, we then cross-correlated each individual burst with the mean burst and extracted the maximum cross correlation (from the centre point ± 25 ms; see methods) and averaged this across all bursts. This demonstrated a mean maximum cross-correlation of 0.39 ± 0.03 and 0.44 ± 0.03 for the ERD and ERS periods respectively (Fig 5B).

**Figure 5.**
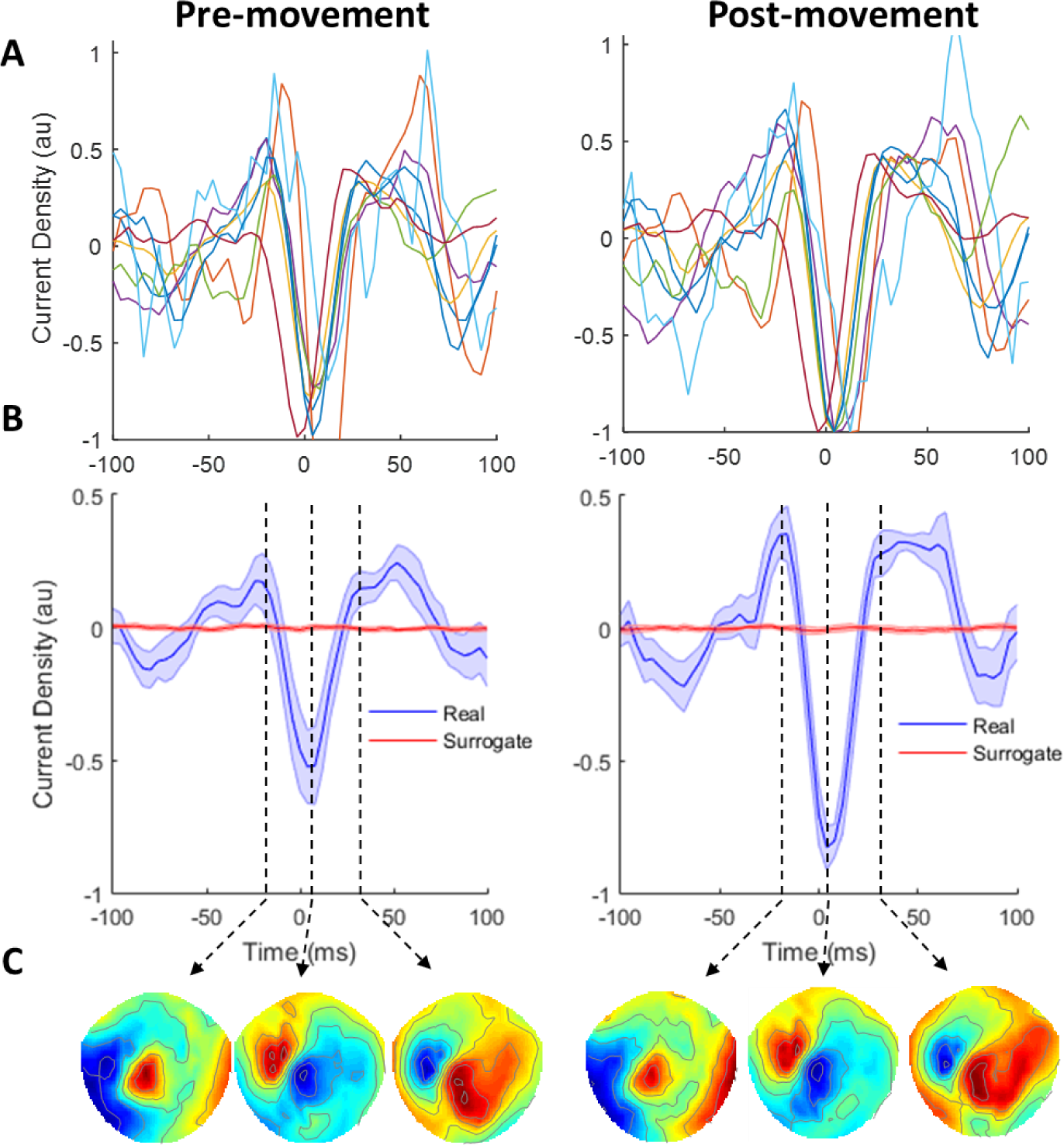
Beta bursts pre- and post-movement in source and sensor space. **A** Individual (mean and normalised) bursts from all subjects shown in the time domain, source (M1) space in the pre- (left) and post- (right) movement periods, respectively. **B** Mean (normalised to average ERS burst max) beta burst across all subjects for real (blue) versus phase shuffled, surrogate (x1000 shuffles), data (red), shown separately for the pre- (left) and post- (right) movement periods. **C** Sensor level scalp maps showing sensor level activity (normalised within subjects and averaged over all bursts / subjects) aligned to three periods of maximum extrema (indicated by vertical dashed lines).

The subject specific mean burst shapes (real data and surrogate) were then taken forwards for analysis of between subject effects (normalised by the maximum extrema). These highlighted a relatively conserved mean burst shape across the pre- and post-movement periods, albeit with larger bursts post-movement, across subjects (Fig 5B), a finding which was not found in the surrogate data. This suggests that although there is a stereotyped average beta burst shape across subjects, there is a significant degree of variability across bursts within subjects. Finally, in order to determine if the source reconstructed beta signal corresponded to a single dipole, we analysed the sensor level data using the same epochs as source level data at a single sensor (MLT14) overlying the left sensorimotor cortex. This demonstrates a single dipole at the maxima of the beta burst (4ms) which is highly stereotyped in both the pre- and post-movement bursts (Fig 5C).

### Spatial distribution of beta bursts and conventional ERD and ERS

Having demonstrated that beta bursts are transient, we proceeded to examine the distribution and consistency of beta bursts over space. Previously, movement related *average* beta changes have been considered to be spatially diffuse and bilateral in keeping with a global state parameter such as (new) movement inhibition (Doyle et al., 2005; Engel and Fries, 2010; van Wijk et al., 2009, 2012). However, we considered whether the spatially diffuse signal may be partially a result of averaging of bursts. We therefore analysed the spatial topography of the ERD and ERS movement-related beta change (dB normalised to −3000 → −2500 ms pre-movement, shown in classical top panel spectrograms) during the pre- and post-movement periods for each subject in source space. During the pre-movement (ERD) period (Figure 6C: left panels), the reduction in average beta power was spatially distributed and bilateral in the majority of subjects, echoing previous reports (Cheyne, 2013; Doyle et al., 2005). This spatial pattern was conserved in the post-movement (ERS) period although there was some notable heterogeneity across subjects (Fig 6C; right panels).

**Figure 6.**
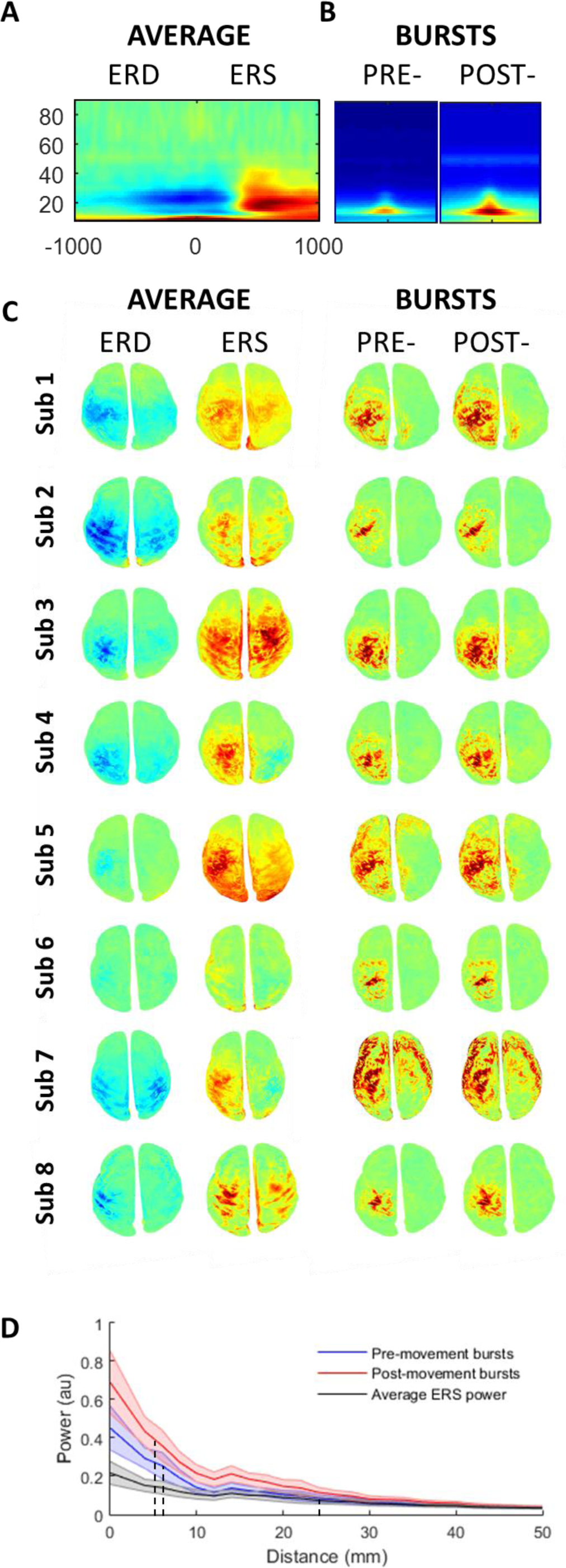
Topography of classical average beta ERD and ERS signals is more spatially diffuse than beta bursts. **A** Conventional time frequency spectrogram (baseline (dB) normalised power) averaged across all subjects at M1 contralateral to movement (top panel). **B** Time frequency spectrogram (same timescale as conventional analysis −400 → 400 ms) aligned and averaged over all bursts and subjects for the pre- and post-movement periods. **C** Spatial topography for each individual subject during the ERD period (left panel; −200 ms → 0 ms) and ERS period (right panel; 400 ms → 800 ms), normalised to baseline (dB, −3000: −2500 ms, left two columns). This is contrasted against the spatial topography of the individual bursts during the same periods (right two columns) normalised to the beginning of each burst (dB; 100 ms prior to amplitude peak). Note that the bursts have a similar pattern but more focal topography than the ERD and ERS (see supplemental video 1 & 2). **D** Topography of the drop off in activity from the M1 cortical area for all subjects ± SEM (normalised within subject), for the ERS as well as for the pre- and post-movement bursts. This demonstrates a much faster drop off in activity (with distance from M1) for the bursts (FWHM: ERD 7.25 ± 0.8 mm, ERS 6.75 ± 0.75 mm) than for the classical average ERS (FWHM: 25mm).

The spatial topography of the ERD and ERS was then directly compared with the spatial topography of the average beta burst. Source level beta burst topography (dB; normalised to beginning (100 ms pre-peak)) was averaged across all pre- and post-movement bursts separately to generate a topographical map of the spatial distribution analogous to the ERD and ERS maps (Figure 6B; right panels). This demonstrated a similar topography of bursts between pre - and post-movement epochs, although post-movement bursts contained slightly greater power. Direct comparison of the spatial topography of the beta burst and the average ERD and ERS revealed that the topography of the bursts was overall more focal than that found for the ERD / ERS spectral changes. We formally quantified this by examining the reduction in normalised power as a function of distance from the primary motor cortex, for the ERS peak and for individual bursts (separately for pre- and post-movement bursts). While the peak height of the bursts was higher compared to the maximum ERS, the decrease in activity with distance away from the centre of the primary motor cortex was significantly steeper for bursts compared to the ERS period beta activity (Fig 5C). For the pre- and post-movement bursts, the Full Width Half Maximum (FWHM) was, on average, 7.25 ± 0.8 mm and 6.75 ± 0.75 mm, respectively. By contrast, for the conventional trial averaged peak ERS, the FWHM was significantly greater (more diffuse) at 25 ± 2.4 mm (t_7_=7.4, p<0.001 and t_7_=7.8, p<0.001). Therefore, we here show that beta bursts are, spatially more focal than conventional trial averaged ERS activity.

### Beta burst timing predicts behaviour

Having shown that conventional, pre-movement beta ERD and post-movement ERS are dominated by punctate, non-rhythmic, spatially focal, high amplitude burst events, we proceeded to examine the putative functional relevance of these beta burst events in the cued movement selection task. All subjects were able to complete the task and mean response time was 277.5 ± 29.6 ms with a mean error rate of 12.6 ± 4.7 % (Fig 7B).

**Figure 7.**
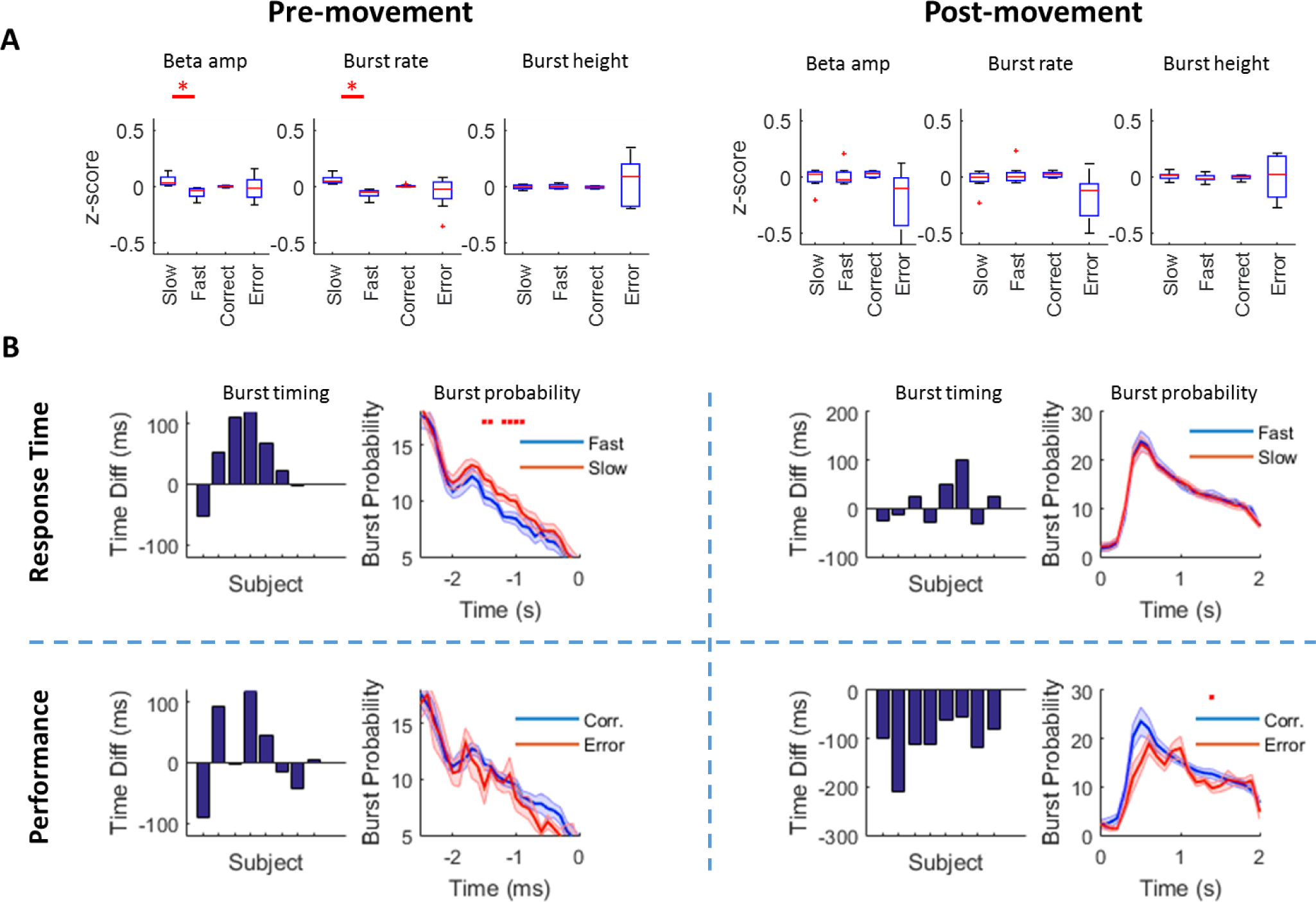
Effect of beta amplitude, burst rate and burst timing on response time and performance. **A** Relationship between z-scored beta amplitude, burst rate and burst height for slow versus fast trials and correct versus incorrect trials in the pre- (left) and post- (right) movement periods respectively (FDR corrected significant tests denoted by red asterisk). Note that in the pre-movement period, the beta amplitude and burst rate are predictive of response time (but not errors) whereas in the post-movement period the relationship is between burst rate and movement errors (although this does not survive FDR correction; p_FDR_=0.06). **B** Relative burst timing (bar chart) and burst probability (line graph) in the pre- (left) and post- (right) movement period for response time (upper panels) and errors (lower panels). Bar charts show the absolute difference in the timing of final burst prior to the instruction cue (slow - fast trials and correct - error trials) pre-movement and the absolute difference in the timing of the first burst (slow - fast trials and correct - error trials) in the post-movement period. This demonstrates that pre-movement, the timing of the final burst is later in slow trials in 6 out of 8 subjects and post-movement the timing of the first burst is earlier in 8 out 8 subjects on correct trials. Burst probability plots show the window over which this burst timing operates and demonstrates a higher rate of bursts (ie higher likelihood of a burst in this window on any given trial) in slow versus faster trials between 1500 → 500 ms (red asterisks, FDR corrected). Conversely, post-movement, burst probability plots shows a significant (FDR corrected, red asterisk) difference between burst probability in correct (blue) versus incorrect (red) trials across all subjects (~1500ms, lower panels). The difference at the first peak (600 ms) did not survive FDR-correction.

It has previously been demonstrated that pre-movement beta activity relates to response time (Doyle et al., 2005) and post-movement beta relates to error (Tan et al., 2014, 2016; Torrecillos et al., 2015). As we observed that the ERD and ERS are tightly coupled to beta burst rate, we predicted that beta burst rate in the pre- and post-movement periods would relate to response time and error monitoring, respectively.

We therefore divided trials according to response time (median split; slow versus fast trials), and compared the z-scored mean (averaged over the time period −2500 → 0 ms pre-instruction cue and averaged over fast or slow trials separately) beta amplitude and burst rate in the pre-movement period in these two groups of trials for all subjects. Pre-movement mean beta amplitude distinguished between fast and slow trials (t_7_=3.2, p=0.015, p_FDR_=0.015), with faster trials displaying, on average, lower beta amplitude (Fig 7A). We then examined beta burst rate over the same period with a matched analysis that showed that beta burst rate was also lower on trials with faster compared to slower response times (t_7_=4.5, p=0.003, p_FDR_=0.006).

We next turned to the post-movement period (0-2000 ms) to investigate the relationship between beta amplitude, burst activity and the commission of errors. We therefore divided trials into correct versus incorrect trials and compared z-scored mean (averaged over the time period 0 → 2000 ms post movement and averaged over correct and incorrect trials separately) beta amplitude and burst characteristics in these two groups of trials for all subjects. Trials with errors were weakly related to post-movement mean beta amplitude (t_7_=2.1, p=0.07, p_FDR_=0.07) and burst rate (t_7_=2.7, p=0.03, p_FDR_=0.06), whereby errors were associated with lower average beta amplitude and burst rates, although this did not survive FDR multiple comparison correction.

In order to check whether the pre- and post-movement beta amplitude and burst rate were specific to response time and error respectively we performed secondary (reversed) analyses. We asked whether pre-movement beta related to decision error or post-movement beta related to response time but found no significant effects (p>0.1 for all correlations, uncorrected). By way of further control analyses, we also investigated other beta burst features and found no relationship between pre-movement beta burst height (t_7_=0.29, p=0.78) or duration (t_7_=0.37, p=0.73) and response time, nor with post-movement beta burst height (t_7_=0.03, p=0.9) or duration (t_7_=0.01, p=0.99) with errors. Finally, beta amplitude and burst rate in the surrogate phase shuffled dataset did not correlate with response time or commission of error in either pre- or post-movement periods (p>0.05 all tests; uncorrected). These results pinpoint a functional dissociation between pre-movement beta (response time) and post-movement beta (performance error).

However, both mean beta amplitude and beta burst rate (across trials) conceal whether the precise time of occurrence of beta bursts is behaviourally relevant, which would be expected if they contribute to direct communication (Palmigiano et al., 2017) or local processing (Spitzer and Haegens, 2017). Moreover, there is behavioural evidence in the sensory cortex linking beta burst timing to sensory thresholds (Shin et al., 2017), although demonstration for this in the motor system in humans is lacking. We predicted that beta burst timing prior to movement would relate to response time, and beta burst timing after movement would be associated with the commission of errors. To test this, we first commenced with a conservative analysis and examined mean differences in burst timing across trials with respect to behaviour. For this, we again split trials according to fast versus slow trials for the pre-movement period, and correct versus incorrect trials for the post-movement period.

The mean timing of the final beta burst before movement onset was earlier on fast compared to slow trials in 6 out of 8 subjects (mean: 42 ± 22 ms; Fig 7B left panels). We then examined the time course of this difference by plotting the mean burst probability on fast versus slow trials across all subjects and found a critical window between −1.5 s and −0.9 s where the occurrence of bursts during this window predicted slower response times (p<0.05, FDR corrected, Fig 7C upper panels). Burst timing prior to movement onset was not related to movement errors (p>0.05, Fig 7C lower panels).

Post-movement, the timing of the first post-movement beta burst occurred later on error trials compared with correct trials in all 8 subjects (mean: 105 ± 17 ms; Fig 7B right panels). When contrasting burst probability at each time point for correct versus incorrect trials, we observed two periods of lower burst probability for incorrect trials at 600 ms (p = 0.026; p_FDR_ = 0.19) and at 1400 ms (p = 0.0022; p_FDR_ = 0.046, Fig 7D, upper panel), the latter of which survived FDR correction. Post-movement burst timing did not correlate with previous response time (p>0.05, Fig 7D lower panels).

In order to further examine the effect on behaviour of burst timing, on a trial-by-trial basis and also to compare this with beta amplitude and beta burst rate, we performed a regression analysis with linear mixed effects on the z-scored trial-wise beta amplitude, burst rate and burst timing (see methods). For each trial, we determined the timing of the last beta burst prior to the instruction cue, and the first beta burst post-movement for regression against response time and error commission, respectively.

In the pre-movement period we used a linear mixed effects model including trial-wise mean beta amplitude, burst rate and the timing of the final burst in the movement preparation period (− 2500 ms → 0 ms) prior to the instruction cue and their effect on response time. We found that both beta amplitude (χ^2^(1)=5.88, β=0.0031, p=0.015) and beta burst timing prior to instruction cue (χ^2^(1)=6.6, β=0.0022, p=0.010) significantly related to response time, but no such relationship was found for beta burst rate (χ^2^(1)=1.2, β =0.0015, p=0.27) when accounting for the other factors.

In the post-movement period, we used a generalised linear mixed effects model to assess the combined effect of trial-wise beta amplitude, burst rate and the timing of the first burst following button press on error commission (0 → 2000 ms). With all three variables in the model together, only the timing of the first beta burst (W(1)=28.0, β =−0.21, p=1e-07) was significantly related to error, with beta amplitude (W(1)=2.1, β =0.09, p=0.15) and beta burst rate (W(1)=0.001, β =−0.002, p=0.9) showing no significant effect when accounting for all three variables.

Overall, these results demonstrate that the timing of beta bursts closely relates to behaviour on a single trial level. This effect was stronger post-movement and related to errors where it factored out the effect of beta amplitude even at the single trial level. This supports the contention that behaviourally, bursts and their timing, not average (both within and across trials) amplitude per se, might be key to understanding the functional relevance of beta and its relationship to the motor system.

## DISCUSSION

Movement-related activity changes in the beta frequency band (13-30 Hz) are a hallmark feature of healthy and pathological movement, yet their functional role remains highly debated (Jenkinson and Brown, 2011; Kilavik et al., 2012; Spitzer and Haegens, 2017; van Wijk et al., 2012). Overcoming this knowledge gap is critical to understanding the physiological underpinnings of healthy and pathological movement, and to inform the development of new treatments for diseases of movement that show pathophysiological beta activity (Crowell et al., 2012; Hammond et al., 2007; Little et al., 2013).

### Average beta amplitude analyses conceal burst timing dynamics

It is increasingly becoming apparent that beta amplitude and particularly average beta changes in the sensorimotor system may in fact be a simplistic summary of the underlying neural dynamics (van Ede et al., 2018; Feingold et al., 2015; Shin et al., 2017). Further understanding of the role of these signals mandates detailed decomposition of the average changes of rhythmic activity into the underlying dynamics at a single trial level. By leveraging high-precision MEG, we here show that activity at the single trial level is highly dynamic, and is dominated by bursts of beta activity that dictate the majority of changes in synchronization and de-synchronization classically seen (on average) before and after movement, respectively. This is despite the fact that beta bursts comprise a relatively small fraction of total time in a given trial (pre-movement 11.9 %; post-movement 12.7 %), with a mean burst duration of ~100ms, and burst rate of ~1.5 bps, even during the sustained average amplitude increases generally seen post-movement (burst < 1 in 4 trials at ERS peak). Importantly however, this burst activity is modulated by the motor response suggesting a behaviourally relevant role that is not well captured in the average beta amplitude alone. Specifically, the timing of the beta burst before and after movement predicts behaviour, and notably, in the post-movement period, this factored out the effect of beta amplitude. In the pre-movement period, we found that bursts near to the instruction cue were followed by slower response times. By contrast, in the post-movement period, beta bursts occurred earlier when a correct response was made. This timing relationship with behaviour is concealed in conventional analyses of trial averaged beta amplitude.

### Beta burst timing in movement planning and performance monitoring

Our results provide a novel link between cortical beta burst activity in the primary motor cortex to endogenous movement planning and performance in humans. This extends recent work on the short lasting temporal dynamics of beta in the human sensory cortex, basal ganglia and non-human motor system (Leventhal et al., 2012; Little et al., 2012b; Sherman et al., 2016; Shin et al., 2017; Tinkhauser et al., 2017a). These findings collectively support a picture of beta being mechanistically relevant (Androulidakis et al., 2006, 2007; Gilbertson et al., 2005; Joundi et al., 2012; Lalo et al., 2007) and characterised by bursts which are temporally scattered and sparse, with beta burst rate and timing predicting motor behaviour, as shown here. Our results motivate a reappraisal of the divergent class of interpretations of beta, including processes relating to status quo maintenance, motor attention, motor idling, simple movement preparation or inhibitory signalling, and which have been generally dependent on trial averaged analyses (Donoghue et al., 1998; Engel and Fries, 2010; Pfurtscheller, 1981; Pfurtscheller et al., 1996; Sanes and Donoghue, 1993).

### Beta bursts suggest active processing

Our results here suggest that, rather than beta activity sub-serving a generalised merely inhibitory role, it has a specific, information encoding role that can rapidly bias behaviour on a trial-by-trial basis despite the events being short lasting. Such a view sides with recent proposals that link beta activity to more complex information coding, such as response uncertainty (Grent-’t-Jong et al., 2014; Rhodes et al., 2018; Tzagarakis et al., 2010), biasing motor response outputs to mediate affordance competition (Grent-’t-Jong et al., 2013, 2014; van Wijk et al., 2009), or even extensive top down hierarchical cortical processing (Bastos et al., 2012; Fries, 2015; Haegens et al., 2011; Palmer et al., 2016; Pape and Siegel, 2016). Here we demonstrate the rapidly dynamic nature of movement-related beta bursting that could be now incorporated into future modelling and theorising of beta in the motor system and potentially more widely.

What informational coding role might these bursts relate to? Higher powered beta bursts at the cellular and network level must signify synchronous recruitment of large neuronal populations to fire together over just a few cycles (Buzsáki and Freeman, 2015). This could potentially relate to short lasting activations of local cortical representations with an active, task specific, information carrying role with beta controlling the balance of local encoding versus onward transmission (Brittain et al., 2014; Naud and Sprekeler, 2018; Palmigiano et al., 2017; Sanes and Donoghue, 1993; Spitzer and Haegens, 2017; Swann et al., 2009). This is supported by our finding that burst activity was spatially more restricted and more transient than previously appreciated and by previous studies showing improved corrective responses during periods of high beta (Androulidakis et al., 2006, 2007; Gilbertson et al., 2005). If beta bursts do indeed relate to the transient activation of cortical representations how might this relate to the planning and execution of movement? Our results here support the recruitment of similar neuronal populations in the pre- and post-movement periods through the demonstration of highly similar patterns of activity before and after movement in the time domain, source space and on sensor maps (albeit fractionally larger post-movement). This similarity is despite greatly differing behavioural demands during these periods. Beta bursts prior to movement could then reflect the orchestration of incoming information for the formation of action goals. Once a goal is specified, no further rehearsal is required and bursts effectively stop as the moment of movement approaches.

By contrast, in the ERS period following movement, the updating of action goals based on the outcome of a movement is required for learning, and a prominent role for beta in this process is consistent with the idea that beta activity tracks a history of inputs and errors (Gould et al., 2011, 2012; Tan et al., 2016; Tzagarakis et al., 2010). One could also speculate that the delay seen in the beta burst following errors allows for more information gathering and sensori-motor integration. Others have suggested that beta bursts may be simply represent inhibitory processing, even if transient and local (Sherman et al., 2016; Shin et al., 2017). Although, we here provide indirect evidence for an active role, full arbitration will be dependent on causal interventions, ideally spatiotemporally focused stimulation which are currently limited (Joundi et al., 2012; Lalo et al., 2007; Nowak et al., 2017).

Notably, subcortical beta bursting is now the basis for developing adaptive deep brain stimulation (DBS) technologies that seek to directly reshape the temporal profile of beta events and thereby provide evidence for its causal role (Little and Bestmann, 2015; Little et al., 2013, 2016, Tinkhauser et al., 2017b, 2017a). If effective beta burst triggered adaptive DBS could be demonstrated at the cortical level, this will not only validate the causal, mechanistic effects of beta bursts on behaviour but would also open up treatment possibilities for neurological disorders characterised by aberrant cortical beta activity (de Hemptinne et al., 2013; Swann et al., 2018). Interestingly, emerging evidence suggests that other frequency bands may be also be better characterised by bursting rather than sustained oscillations (Fig S3), in particular gamma and theta frequency activity (Chandran Ks et al., 2017; Lundqvist et al., 2016; Zavala et al., 2016). Whether short lasting oscillatory bursts over the full range of frequencies represent a widespread mechanism of endogenous, local, content reactivation (Spitzer and Haegens, 2017) or can also support long range communication (Fries, 2015) more generally remains to be determined.

Here we specifically focused our analysis on beta burst rate and timing, and demonstrate that these can account for the changes seen in the ERD and ERS both in terms of the neural signatures (slow changes in beta amplitude are, in the main, the summation of bursts) and behaviour (burst timing is strongly related to motor behaviour and for errors this relationship is stronger than that found with conventional beta amplitude, even at the single trial level). Notably, if beta bursts do represent active processing, then further information will likely be present in the time series shape that is lost in conventional spectral analyses (which effectively enforce a frequency specific sinusoid), and which is becoming accessible through emerging techniques (Cole and Voytek, 2018; Fransen et al., 2015; Whitten et al., 2011).

### Spatial Topography of Beta Bursts revealed by high resolution MEG

With regards to a putative functional role, it is also worth noting that the spatial extent of classical beta changes (ERD and ERS) is often topographically distributed and bilateral. Interestingly, using a matched, parallel analysis on beta bursts, these appear more focally consistent than previously appreciated. Our topographical maps are of the *average* (classical) ERD and ERS and *average* bursts. Indeed we would expect that the trial-wise, spatial network, beta burst topographical configurations will likely be more complex and dynamic than shown here (including for the *average* beta burst) and may also contain important and behaviourally relevant information that may traverse the cortex over time. Methods for dealing with this level of spatio-temporal complexity such as Independent Component Analysis and Hidden Markov Modelling are now validated and will in the future enable the community to address whether the time varying spatial topography of beta burst (Baker et al., 2014; Lee et al., 2003), or their propagation over the cortex (Rubino et al., 2006) also contribute to behaviour in the same way as burst timing does, as shown here.

## Conclusion

Beta activity in the primary motor cortex in humans is primarily characterised by brief, high amplitude, sporadic bursts of activity. These bursts are spatially more focal than previously shown and through summation account for the majority of classical, trial-averaged, ERD and ERS signals. Moreover, beta burst timing is strongly related to motor behaviour with pre-movement beta bursts being associated with delayed movement initiation and early post-movement bursts to correct performance. These results challenge classical interpretations of beta activity that propose a global inhibitory role and suggest a specific informational carrying role of beta burst activity during action preparation and decision monitoring.

## Methods

Eight healthy subjects participated, following informed written consent which was approved by the UCL Research Ethics Committee (reference number 5833/001). All subjects were right handed, had normal or corrected-to-normal vision, and had no history of neurological or psychiatric disease (6 male, aged 28.5 ± 8.52 years).

### Visually cued movement selection task

Participants performed a visually cued movement selection task, in which they were had to press a corresponding (left or right) button as quickly and as accurately as possible after a visual instruction cue (or up to a maximum of 1s in the absence of a response) (**Figure 1**B). The instruction stimuli were preceded by a preparation cue that in 70% of trials predicted the direction of the upcoming imperative stimulus, thus allowing participants to prepare their response in advance of the subsequent instruction cue.

At trial onset, participants fixated on a white cross (0.5° × 0.5°) in the middle of the screen. Following a short delay (uniform random distribution between 1 and 2s), a random dot kinetogram (RDK) appeared with coherent motion to either the left or the right that indicated the likely direction of the upcoming instruction cue (70% of the time). The RDK comprised a 10°×10° square aperture, in the centre of the screen with 100, 0.3° diameter dots, all traversing at 4°/s. There were three levels of coherent motion, which defined the percentage of dots that moved coherently, with the rest of the dots moving in random directions. The middle level of coherence was set individually for each subject using 40 perceptual decision-making trials at the beginning of the experiment, with an adaptive staircase design to achieve 82% accuracy (QUEST; Watson and Pelli, 1983). The lower and higher levels of coherence were then set at 50% and 150% respectively and coherence levels were balanced across trials. The RDK lasted 2s and was followed by a 500 ms delay and then by an instruction cue consisting of a 3° × 1° arrow pointing to the left or the right (Figure 1A). Responses were indicated via a right-hand index (left instruction cue) or right-hand middle finger (right instruction cue) on a button box. The paradigm was implemented using the Cogent 2000 toolbox through MATLAB (The MathWorks, Inc., Natick, MA. http://www.vislab.ucl.ac.uk/cogent.php).

### Magnetic resonance imaging

Subjects underwent both a standard MRI scan for individualised head cast creation and a high resolution, quantitative, multi-parameter map scan (Lutti et al., 2014), which was used for accurate cortical surface extraction and source reconstruction. The first anatomical sequence used a 12-channel head coil and was radiofrequency (RF) and gradient spoiled T1 weighted, 3D fast low angle shot (FLASH) sequence with an image resolution of 1mm^3^, field-of-view: 256,256 and 192 mm along the phase (A–P), read (H–F), and partition (R–L) (3T Magnetom TIM Trio, Siemens Healthcare, Erlangen, Germany). Excitation flip angle was set to 12° to ensure sufficient signal-to-noise ratio and repetition time was set to 7.96 ms. MPM protocol images consisted of 3 spoiled multi-echo 3D FLASH acquisitions (800 µm isotropic resolution, proton density, T1 or MT-weighting) using a 32 channel headcoil with two additional sequences used for calibration and correction of inhomogeneities in the radio-frequency transmit field. To achieve an MT-weighting, a Gaussian RF pulse 2 kHz of resonance with 4 ms duration and nominal flip angle of 220° was applied. The field of view was set to 224, 256 and 179 mm along the phase(A-P), read (H-F) and partition (R-L) directions respectively. Alternating readout gradient polarity (Gradient Echoes) were acquired at eight equidistant echo times ranging from 2.34 to 18.44 ms, in 2.30 ms steps (bandwidth of 488 Hz/pixel, 6 echoes for repetition time of 25 ms for all FLASH volumes). Transmit field inhomogeneities were mapped using a 2D STEAM approach, including corrections for geometric distortions of EPI data (Lutti et al., 2014). Total acquisition scanning time for all protocols was < 30 min.

### Head-cast fabrication

In order to record beta activity with a maximal level of spatial precision we used subject specific head-casts which have been shown to optimise co-registration, reduce head-movements and thereby significantly improve signal-to-noise (SNR) (Meyer et al., 2017; Troebinger et al., 2014a). Anatomical MRI sequences were used to extract scalp surfaces (SPM12) by registering MRI images to a tissue probability map which classified voxels according to tissue makeup (e.g. skull, skin, grey matter etc.). These were then transformed into a surface using the isosurface function in MATLAB (version 2015A, Mathworks, MA, USA). This surface was converted to a standard template library (STL) format with digital outlines of three fiducial coils placed at conventional sites (left / right pre-auricular and nasion). From this digital image, a positive head model was 3D printed (Zcorp, 600 × 540 dots per inch resolution) and this model placed inside a replica dewar, as in our previous studies (Meyer et al., 2017; Troebinger et al., 2014a). Liquid resin was then poured between the two surfaces, resulting in a flexible, subject specific head-cast (Meyer et al., 2017).

### MEG scanning

Subjects underwent MEG recordings (CTF 275 Omega system) using their individual head-cast during a visually cued, movement selection task. Head position was first localised using the three fiducials placed at the nasion and left/right pre-auricular points, within the headcast. Data was sampled at 1200 Hz before being imported to SPM 12, and subsequently downsampled to 250 Hz. One advantage of the head-cast approach is that it permits multiple recording sessions within each subject with the head being in the same position. This allows for recording large scale individual datasets, but because of the lack of movement and low co-registration error, this does not result in accumulative SNR decreases (Meyer et al., 2017; Troebinger et al., 2014a). Movement of the fiducials was continuously measured by the CTF system and the standard deviation of movement averaged across the 3 fiducials and presented separately for the x, y and z directions (Fig 1C). Subjects performed 1 - 4 recording sessions, with each session split into 3 separate runs. Participants completed 3 blocks per session (with short breaks between them), and 1-4 successful sessions on different days. This led to a total number of 1614 ± 763 (mean ± sd) completed trials per subject taken forward for analysis. Data from individual runs within sessions were concatenated and these datasets from each session analysed individually and results averaged across session for each subject. The data was filtered with a 5th order (Butterworth) bandpass filter (2- 100 Hz, Notch filter: 50Hz) and downsampled to 250 Hz. Eye blink artefacts were removed using multiple source eye correction (Berg and Scherg, 1994) and trials with variance > 2.5 sd away from the mean were excluded. Data was epoched from 3.5 s before to 1.5 s after the instruction cue (total 5 s) for the analysis of pre-movement neural activity (cue-locked) and to 2 s before to 2 s after the button press for post-movement analysis (response locked).

### Source inversion

For source analysis, sensor level data were inverted onto subjects’ individual cortical surface meshes. These meshes were extracted using Freesurfer (v5.3.0; Fischl, 2012) from multi-parameter map using the PD and T1 maps as inputs with in house modifications of the reconstruction to avoid tissue boundary segmentation failures (Carey et al., 2017). Each mesh was then down-sampled by a factor x 10 (~33,000 vertices per mesh) and smoothed (5mm) (Carey et al., 2017).

Source reconstruction (estimation of current dipole positions and strengths) was performed using SPM12 and an Empirical Bayesian beamformer (with a supplemental analysis for comparison by Minimum norm inversion; Figure S2) without Hanning windowing, using a Nolte single shell model and a frequency of interest of 1 – 90 Hz (Belardinelli et al., 2012; López et al., 2014; Nolte, 2003). Data were extracted at each individual vertex as a virtual electrode by multiplying the sensor level data by the weighting matrix (M) between sensors and source from the inversion and the data reduction matrix (U) that specifies the significance of the modes of data that map to the cortex. This estimated time series data was then taken forward for the analysis of beta bursts by selecting the time series from individual vertices closest to the primary motor (M1) cortex which was visually identified for each individual subject with reference to the hand knob (Dechent and Frahm, 2003; Yousry et al., 1997).

### Time frequency decomposition

In order to obtain the beta amplitude trace, the time series virtual electrode data was filtered using a 4^th^ order (two pass) Butterworth filter with a frequency range of 13 – 30 Hz. A complex (analytic) function was then extracted from the time series using the Hilbert transform and the amplitude derived by taking the modulus of that signal:

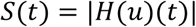

Where S(t) is the amplitude signal, u(t) is the filtered data and H represents the Hilbert transform. For time resolved spectrograms (Fig 2A and Fig 6A/B), a Wavelet convolution was implemented (ft_specest_wavelet script in Fieldtrip - Morlet Wavelet, width = 10, gwidth = 5, (Oostenveld et al., 2011)).

### Identification of beta bursts through empirically defined thresholds

In order to identify beta bursts, we first needed to select an appropriate threshold. Previous studies have generally selected heuristic thresholds relative to the mean beta amplitude and shown robustness of findings across a range of threshold levels (Feingold et al., 2015; Tinkhauser et al., 2017b, 2017a). This works well for individual studies but limits the scope for comparison across different brain sites and recording methods as well as theoretically subsampling a smaller distribution of relevant bursts (if the threshold is set too high), or alternatively, including smaller, non-burst, activity such as noise which may dilute behavioural effects. Recently an empirical method has been introduced whereby burst frequency is correlated with total beta power in the trial across a range of thresholds and the maxima in the correlation used to define the threshold for defining bursts (Shin et al., 2017). Here we used a similar technique to that previously described but defined the threshold in terms of standard deviations away from the median signal (as opposed to multiples of the median) and so as not to transform the underlying signal in any way performed this on the beta amplitude, rather than the beta power (amplitude.^2).

Therefore, for each subject we correlated the trial-wise mean beta amplitude with the number of burst events in each trial (defined according to an amplitude threshold), separately for pre- and post-movement periods. Each subject performed 540 trials per recording session. Following automated rejection, this may leave, for example, 535 valid trials. In such a case therefore, we would correlate the 535 mean amplitudes (across 13 – 30 Hz) in the pre-movement period against the 535 burst counts in the same pre-movement period (repeated separately for the post- movement period). This procedure resulted in one correlation value for each subject for the pre-movement period and one correlation value for the post-movement period, per burst definition threshold. This was then repeated across a range of thresholds (defined in terms of SDs above the median beta amplitude) to generate two curves for each subject, which showed the relationship between amplitude thresholds used to define the burst and the correlation coefficient between the beta amplitude and burst counts across trials (Shin et al., 2017). Curves were then averaged across all subjects and the peak taken at the group level (Fig S1), as the definition of the amplitude thresholds for defining bursts. This revealed a peak correlation between average beta power and burst amplitude at 1.75 standard deviations above the median, which was consistent for both the pre-movement, cue–locked period, and the post-movement, button-locked period (Fig 2C). The threshold of 1.75 SDs above the median was then used for each individual subject, defined according to their own dataset, so that they had statistically matched thresholds, although the absolute threshold levels could differ to take account of varying SNR across subjects. Notably, this threshold was robust to MEG inversion method (Fig S2).

Comparing our empirically derived threshold with previously used burst definition thresholds, we find that this is higher than has been used in subcortical recordings (Tinkhauser et al., 2017b, 2017a), is comparable to findings in the motor cortex in primates (Feingold et al., 2015), but lower than shown previously in the human sensory cortex, (Shin et al., 2017). The fact that this threshold level found here is lower than that of other MEG data may partly related to the use of amplitude correlations rather than power correlations (Shin et al., 2017). Indeed, using power (as opposed to amplitude) increases the empirically derived threshold, but also leads to flatter correlation curves in our dataset (Fig S1). Furthermore, the difference in threshold may likely reflect the difference in recording technique used – namely head-cast MEG which results in higher SNR datasets which one would expect to lead to lower thresholds through reductions in noise (Meyer et al., 2017; Troebinger et al., 2014b, 2014a). Having empirically defined our threshold (x 1.75 sds > median amplitude), bursts were identified by locating peaks above this threshold in the beta filtered amplitude traces from the M1 source localised data and the distribution of these peaks analysed with respect to behaviour. The 1.75 sd threshold (threshold 1) was used to define burst occurrence, with the edges (skirt) of the bursts (threshold 2) was defined as being 1 sd above the median so as not to underestimate the burst length by just examining the central peaks (Shin et al., 2017).

### Beta burst timing

Burst probability (at each time point *across* trials) was calculated by binary coding all trial time points as to whether the beta envelope was either above or below threshold 2 on each individual trial and this matrix (trials x time points) averaged over trials and scaled by total trial number to give a time resolved probability of bursts per subject. Time resolved ERD and ERS were calculated by averaging beta amplitude across trials and this average value was then normalised as a % change to baseline in order to facilitate comparison across all subjects (Fig 3A). With the purpose of quantifying the impact of removal of bursts on the conventional ERD and ERS measures, bursts were identified (above threshold 1) and replaced by the trial averaged mean beta amplitude for the pre- and post-movement periods separately (to the edge of the burst - above threshold 2). Individual beta amplitude traces (with bursts replaced) were then averaged across trials to demonstrate the effect of burst removal on the ERD and ERS separately (Fig 3A, red trace = ERD and ERS with bursts replaced). The ERD and ERS (over time) was then correlated against the burst probability curve (over time) to derive an R^2^ value of correlation between the two and plotted as a scatter plot (with a line of best fit) for the pre- and post-movement periods separately. This is a measure of how the shape of the time resolved burst probability (Fig 3A; black trace) matches that of ERD and ERD (Fig 3A; red trace) during the pre- and post-movement periods respectively. Burst amplitude peak timings were also plotted in the form of a raster plot across all trials for all subjects to show the change in burst timing with regard to behaviour (Fig 3C).

### Beta burst characteristics

We first analysed beta amplitude (all beta amplitude, not restricted to bursts) variability and quantified this by the coefficient of variation (CV = std beta amplitude / mean beta amplitude). This was performed for each trial for the pre- and post-movement periods separately and plotted as a histogram of CV values (across trials) for a single subject (Fig 2A) and for all subjects together (Fig 4A). This was then compared to a spectrally matched surrogate dataset that retained all of the spectral features of the original data except, through phase shuffling (prior to filtering), the time series was randomised and therefore redistributed the (same) beta power across time, taking a highly non – stationary signal and making it more stationary (function: spm_phase_shuffle; SPM12) (Feingold et al., 2015). Having established that the beta amplitude is more variable than would be expected from a spectrally matched surrogate dataset, we proceeded to examine the beta bursts specifically. Burst (as defined by beta amplitude periods above our empirically defined, 1.75 sd > median, threshold) characteristics were quantified by height and duration. Height was taken as the maximum of the peak of the burst in the amplitude domain. Duration was defined as the time (ms) that the amplitude remained above threshold 2. These features were again compared to surrogate data using the phase shuffled dataset as described. The differences (real data – surrogate data) in the burst characteristics between the two different populations (same threshold used for burst definition in both datasets) were plotted for each subject on a 3D histogram. We then tested the hypothesis that beta bursts were larger than would be expected by comparing the distributions of burst heights and burst lengths for 10 different heights (percentiles) and lengths (100 ms bins up to 1000 ms) against our surrogate, spectrally matched distributions (mean of x1000 iterations) using paired t tests across subjects (FDR corrected). This allowed for ruling out the possibility that beta burst activity was not just noise superimposed on top of a slowly changing signal, which would predict a similar distribution of CV and beta bursts in a surrogate dataset.

To analyse the shape and consistency of the bursts we took our data and re-aligned it to the centre of the peak of the amplitude. Unfiltered source level data was re-epoched around the amplitude peaks with a window of 100 ms on either side (Fig 5A). This burst aligned (unfiltered) dataset was then averaged across all bursts to look for consistency of the waveform (in the time domain) within the amplitude aligned bursts and this was repeated with surrogate data (Fig 5A). Individual bursts were then cross-correlated (r) against the average burst and the maximum r value (max(r)) within 25 ms (half a cycle), before or after the midpoint (to allow for some a small amount of jitter = ~1 beta cycle), taken as a measure of the concordance of the individual burst with the average, repeated for the pre-movement and post-movement bursts. The average waveform for each subject was then normalised by the maximum (absolute) value for that subject and these normalised subject specific bursts – averaged across all subjects with the time series +/- orientation displayed with the minimum closest to time zero as per previous studies (Sherman et al., 2016; Shin et al., 2017). All statistical analyses were performed on the beta amplitude envelope and were therefore invariant to inherent uncertainty in dipole orientation causes by source inversion uncertainty and sulcal variation across subjects. Analyses were repeated in the theta and gamma ranges as functional controls and to check that the behavioural relationships found with beta bursts were specific (Fig S3).

Finally, the sensor level field map was examined by averaging the (subject mean normalised) magnetic field for all bursts and subjects for a representative sensor overlying the left motor area (MLT14) at the three moments of the average maximum dipole in the amplitude aligned dataset and plotted as an average scalp map which confirmed the presence of a single dipole (spm_eeg_plotScalpData) that was similar for pre- and post-movement periods.

### Analysis of the burst spatial distribution

We examined the distribution of activity on the cortical mesh in source space using both a classical (averaged across trials) relative change (ie ERD and ERS) and a matched analysis for average burst activity changes at the peak. For spatial topography, we used the conventional, baseline corrected, (for comparison across subjects) analysis consisting of averaging the power change of the period of interest (e.g. ERD minimum, ERS maximum, burst maximum) relative to baseline for all different vertices across the mesh:

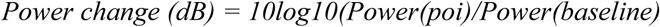

The baseline was defined as the 200ms prior to the onset of the trial, the ERD as the final 200 ms prior to button press and the ERS as 200 ms either side of the time point 600 ms post button press. The power in the bursts was normalised to the start of the bursts (− 100 ms from the peak) using the same formula (dB). The power was then shown projected onto the mesh with colour range limits between – 5 dB and + 5 dB. For the calculation of the FWHM, the power at each vertex was binned according to distance from the centre of the hand knob area (in 5mm bins up to 50 mm) of the primary motor cortex and the power averaged within this bin. This was performed for the ERS power (timing as defined above) and also for each individual burst at the peak (power drop off averaged over all bursts). The average power change by distance for all bursts (pre- and post-movement) as well as the ERS power change by distance were then normalised within subjects by dividing all of them by the maximum power (first bin) of the average post-movement bursts (which was the largest). The FWHM was then calculated within individual subjects as distance away from the centre of the M1 hand area for which the power had dropped by half.

### Behavioural analyses

In order to analyse the differential contribution of the beta amplitude and the burst characteristics with regards to behaviour these features were calculated separately in the pre- and post-movement periods. Responses that matched the direction of the instruction cue were classed as correct and the response time (RT) was taken as the time difference between instruction cue and the button press. Beta amplitude, burst rate, mean burst height and mean burst duration were calculated in the pre-movement period (cue locked dataset; start of RDK −2500 ms → 0 ms instruction cue) and post-movement period (button locked dataset; 0 ms → 2000 ms) and z-scored across all trials. Z-scored trials were then split according to response time (slow versus fast) and decision error (correct versus incorrect) and compared across all subjects by paired t-testing across subjects for both pre- and post-movement datasets respectively. Burst timing was analysed in a similar fashion with the last pre-instruction cue burst timing taken forwards for the pre-movement dataset and also the timing of the first burst after button press taken forwards for the post-movement dataset. Timing resolved burst probabilities across all trials were downsampled to a sampling frequency of 20 and then similarly grouped according to behavioural output (response time and error) with differences compared between trials grouped by behaviour for each time point by paired t-testing (and FDR correction).

Finally, in order to examine the effect of beta amplitude, burst rate and burst timing on a individual trial-by-trial basis and directly compare their impact on behaviour we proceeded to take these features forwards into a linear, mixed effects model (R statistics, version 3.4.3). Firstly, for RT (dependent variable) we used a linear mixed model with congruence, coherence and their interaction as fixed effects and beta amplitude, beta burst rate and beta burst timing plus a subject-specific intercept as random effects. Fixed effects for this model were estimated using type III Wald F tests with Kenward-Rogers approximated degrees of freedom (Kenward and Roger, 1997). For the post-movement analysis and beta amplitude, burst rate and burst timing together we used a generalized linear mixed model with a logit link function, using correct (true or false) in each trial as the dependent variable, congruence (congruent or incongruent) and coherence (low, medium, high) and their interaction as fixed effects, and beta amplitude, burst rate and burst timing plus subject-specific intercepts as random effects. Fixed effects were tested using type III Wald χ^2^ tests.

